# The atypical antipsychotic lurasidone positively modulates the gut microbiota in rats: A comparative study to olanzapine

**DOI:** 10.1101/2024.02.28.582623

**Authors:** Srinivas Kamath, Alexander Hunter, Kate Collins, Anthony Wignall, Paul Joyce

## Abstract

**Background and Purpose:** Antipsychotics like olanzapine are associated with significant metabolic dysfunction, attributable to gut microbiota dysbiosis. A recent notion that most psychotropics are detrimental to the gut microbiota has arisen from consistent findings of metabolic adverse effects. However, unlike olanzapine, the metabolic effects of lurasidone are conflicting, with most reports observing weight loss rather than gain. Thus, this study investigates the contrasting effects of olanzapine and lurasidone on the gut microbiota to explore the hypothesis of “gut neutrality” for lurasidone exposure.

**Experimental Approach:** Using a Sprague-Dawley rat model, the impact of olanzapine and lurasidone administration on the gut microbiota was explored. Faecal and blood samples were collected weekly over a 21-day period to analyse changes to the gut microbiota and related metabolic markers.

**Key Results:** Lurasidone triggered no significant weight gain or metabolic alterations, instead positively modulating gut microbiota through increases in mean OTUs (+50 OTUs) and alpha diversity (+0.5 increase in Shannon’s index). This novel finding suggests an underlying mechanism for lurasidone’s metabolic inertia. In contrast, olanzapine triggered a statistically significant decrease in mean OTUs (−75 OTUs) and substantial compositional variation, suggesting a decrease in microbial richness. Microbiota alterations correlated with metabolic dysfunction, evidenced through a statistically significant 30% increase in weight gain, increase in pro-inflammatory cytokine expression, and increase in blood triglycerides and glycaemic levels.

**Conclusion and Implications:** The study challenges the notion that all antipsychotics disrupt the gut microbiota similarly and highlights the potential benefits of gut positive or neutral antipsychotics like lurasidone in managing metabolic side effects. Further research is warranted to validate these findings in humans to guide personalised pharmacological treatment regimens for schizophrenia.

**Bullet point summary:**

- **What is already known:**

- Olanzapine induces weight gain by disrupting the gut microbiome.
- The impact of lurasidone on the gut microbiome is unknown and weight gaining propensities unclear.
- **What this study adds:**

- Lurasidone positively modulates gut microbiota through enhancement of microbial diversity and richness.
- Potential mechanisms underlying lurasidone’s weight and metabolic neutrality are elucidated.
- **Clinical significance:**

- Gut neutral antipsychotics like lurasidone could be favourable alternatives for patients unable to tolerate olanzapine.
- Personalised treatment for schizophrenia considering individual sensitivities to metabolic effects is emphasised.

## 1. Introduction

Antipsychotics are widely employed for therapeutic management of a range of mental health conditions, including schizophrenia and bipolar disorder. Pathogenesis of these conditions is largely attributed to abnormalities in the synthesis of dopamine, glutamine, GABA, acetylcholine, and serotonin. Justly, antipsychotics used to treat schizophrenia and bipolar disorder demonstrate complex pharmacology and multitargeted modes of action, with impacts on serotonergic, muscarinic, adrenergic, and histaminergic receptors (Ågren, 2021).

Often neglected in mental health pharmacotherapy is the role of the gut microbiota. In particular, the bidirectional interplay between the gut microbiota and mental health, a pathway facilitated by the gut-brain axis and the vagus nerve (Appleton, 2018; Minichino et al., 2023; Radford-Smith et al., 2022; Safadi et al., 2022). The interplay occurs through production and metabolism of gut microbial metabolites, namely short-chain fatty acids and neurotrophic factors which cumulatively lead to regulation of brain-based processes such as regulation of the hypothalamic-pituitary axis, central and autonomic nervous systems (Bear et al., 2021; Clapp et al., 2017; Sarkar et al., 2016, 2020; Spitzer et al., 2021). Prior literature has also associated gut microbiota disturbances with metabolic dysfunction, namely through improper absorption, digestion, and regulation of hormones (Aoun et al., 2020). All these phenomena contribute to dysbiosis associated weight gain.

Recent literature has shifted to investigate the impact of pharmaceutical actives such as antipsychotics and associated excipients on the gut microbiota with many reviews such as one by Minichino et al. concluding their ability to trigger gut microbial imbalances, referred to as dysbiosis (Cussotto et al., 2019; Kamath et al., 2023; Minichino et al., 2023; Subramaniam, Elz, et al., 2023; Subramaniam, Kamath, et al., 2023). For example, Tomizawa et al. analysed 246 stool samples from 40 patients to uncover a dose-dependent generalised decrease in phylogenic, whole-tree and beta diversity indices (all indicative of a dysbiotic gut microbial profile) in patients administered antipsychotics (Tomizawa et al., 2021). Such findings have led to the overall notion that all psychotropics adversely impact the gut microbiota (Ait Chait et al., 2021, 2023; Cussotto et al., 2019; Hao et al., 2023; Minichino et al., 2023; Munawar et al., 2021; Seeman, 2021; Tomizawa et al., 2021).

Further studies such as ones conducted by Qian et al. corroborate the dysbiotic capacity of antipsychotics, namely olanzapine (Qian et al., 2023). The study concluded olanzapine-induced weight gain correlated with longitudinal changes in gut microbiota caused by the drug. Specifically, *Firmicutes* abundance increased from 45% ± 13% to 53% ± 6% (P = 0.0216), with an associated decrease in *Bacteroidota* abundance from 50% ± 14% to 42% ± 7% (P = 0.0320), which is indicative of an obesogenic gut microbial profile (Qian et al., 2023). This, alongside other notable changes (such as decrease in commensal *Akkermansia muciniphila* and *Ruminococcaceae*) suggest potent dysbiosis capabilities of olanzapine. Similarly, Morgan et al. concluded exposure to olanzapine (administered to 8, 6-week-old C57BL/6J mice for 10 weeks at a dose of 50mg/kg) was sufficient to potentiate weight gain (Morgan et al., 2014). Observed gut microbiota findings showcased increase in *Firmicutes* and *Proteobacteria* and decrease in *Bacteroidetes* indicative of a drug-induced obesogenic gut microbial profile. Davey et al. corroborated weight gain tendencies of olanzapine in a Sprague–Dawley rat model with similar observed changes in the gut microbiota, however noted additional variations in metabolic parameters, including increase in adipose tissue, proinflammatory markers (CD68 expression and interleukin (IL)-6 mRNA expression) alongside increased liver weight and carbohydrate regulatory element binding protein (ChREBP) and sterol-regulatory element binding protein 1c (SREBP-1c) mRNA expression (Davey et al., 2013). Plasma pro-inflammatory markers IL-6, -8 and -1β were also elevated.

Lurasidone shares structural and mechanistic similarities to olanzapine. Both possess similar affinities for 5-HT6/7/2A/2C, M1 and H1 receptors located throughout the brain. Furthermore, both drugs possess comparable indications (schizophrenia and bipolar disorder) and comparable adverse drug event profile (including movement disturbances and sedation) (Meltzer et al., 2011). However, despite these similarities, lurasidone’s impact on the gut microbiota remains unexplored and retrospective comparative studies such as one by Pochiero et al. indicate a contrastive weight gain profile compared to most antipsychotics with lurasidone-induced weight gain and weight loss conclusions both being drawn (Pochiero et al., 2021). Thus, lurasidone remains prime for studies exploring its impact on the gut microbiota, serving as the motivation for this study. Here, equivalent doses of lurasidone and olanzapine were administered to Sprague-Dawley rats for a period of three weeks to elucidate the impact of lurasidone on the gut microbiota and metabolic health. The insights derived from this study have the capacity to drastically change future clinical practice relating to prescribed antipsychotic usage by advocating for pharmacotherapies that exert positive or neutral effects on the gut microbiota and associated metabolic health.

## 2. Materials & Methods

### 2.1. In vivo study design

Animal studies were reported in accordance with the ARRIVE guidelines for the accurate and reproducible *in vivo* experimentation and complied with the Principles of Laboratory Animal Care (NIH publication #85-23, revised in 1985) and the National Health and Research Council (Australia) Code of Practice for Animal Care in Research and Training (2014). All animal studies were performed on 8-week-old male Sprague-Dawley rats sourced from Ozgene (Canning Vale, Australia), with 4 female and 4 male rats per group. Rats were housed in a temperature-, humidity- and pressure-controlled animal holding facility with a 12 h/ 12 h light/dark cycle. Rats were randomized upon arrival to the animal holding facility. Researchers were not blinded to the group allocations at any stage of the experimentation or analysis.

Rats were dosed with lurasidone (7.5 mg/kg) or olanzapine (7.5 mg/kg) in PBS (1 mL/kg) via oral gavage for 21 days. A control group was dosed PBS (1 mL/kg) via oral gavage. A longitudinal study was performed to assess the impact of antipsychotic dosing on microbiota changes of individual animals. The sample size for each group (*n* = 8) was based on Power calculations for anticipated changes in Shannon’s Index (microbiota alpha diversity), using a power level of ≥ 0.8 and significance level of 0.05. The study was approved by the South Australian Animal Ethics Committee under approval number U38-22 and U21-23. Rats were housed in groups of two with *ad libitum* access to a normal chow diet and water throughout. Evening dosing (between 16:00 and 18:00) was selected to mimic rodents’ natural behaviour. On Day 22 (i.e. the morning after the final dose), rats were anaesthetized under 5% isoflurane prior to cardiac puncture followed by terminal cervical dislocation.

### 2.3. Gut microbiota analysis

On Day 0, 7, 14, and 21, individual animal faecal pellets were collected during handling and immediately stored at −80°C in sterile tubes to prevent contamination and degradation. Samples were sent to the Australian Genomics Research Facility (Brisbane, Australia) for DNA extraction, PCR amplification, and 16S rRNA sequencing. DNA extraction was performed on one faecal pellet per animal per timepoint to maintain traceability and integrity of microbiota signatures. Subsequently, the 16S rRNA sequences underwent processing for the V3-V4 hypervariable regions, with raw reads clustered into operational taxonomic units (OTUs) using Quantitative Insights into Microbiology Ecology (QIIME 1.8) and the Silva reference database. Taxonomy assignment of OTUs was conducted using the QAIGEN Microbial Insights – Prokaryotic Taxonomy Database (QMI-PTDB) within QAIGEN CLC Genomics Workbench version 23.0.4 (Hilden, Germany). Alpha diversity at the genus level was determined using Shannon’s Index, while beta diversity Principal Coordinate Analyses (PCoAs) were based on Bray-Curtis dissimilarity metrics, both analysed using the QAIGEN genomics module. Statistical significance of microbiota dissimilarities between groups was assessed through Permutational multivariate ANOVA (PERMANOVA) of beta diversity plots.

### 2.4. Metabolic markers

Body weight was monitored twice weekly prior to dosing. Blood samples were collected prior to dose administration on days 0, 7, 14 and 21 before being frozen and stored at −80°C in sterile tubes to prevent contamination and degradation. Blood samples were sent to SA Pathology (Adelaide, Australia) for quantification of blood triglycerides and blood glucose levels.

### 2.4. TNF-α analysis using ELISA

Jejunal tissue samples were promptly frozen post collection in liquid nitrogen for subsequent analysis of, tumour necrosis factor alpha (TNF-α), utilising enzyme-linked immunosorbent assay (ELISA; Invitrogen). The limits of quantification (LOQ) for TNF-α were determined as 11 pg/mL. To prepare the samples, approximately 50 mg of jejunal tissue was weighed and homogenized in 400 μL of RIPA buffer (ThermoFisher Scientific, Australia) containing protease inhibitor cocktail (ThermoFisher Scientific, Australia) at a 1:100 dilution. Protein concentration was determined using a BCA Protein Assay kit (ThermoFisher Scientific, Australia), yielding normalized protein concentrations of 1000 μg/mL in the tissue homogenate. The subsequent steps followed the manufacturer’s instructions, with absorbance readings taken at 450 nm. Cytokine concentrations were then calculated based on the standard curve generated.

### 2.6. Statistical analysis

All statistical analyses (excluding 16S sequencing analyses) of experimental data were performed using GraphPad Prism Version 8.0 (GraphPad Software Inc., California). Statistically significant differences were determined using an unpaired t-test and one-way ANOVA followed by Tukey’s post-test for multiple comparisons. Values are reported as the mean ± standard deviation (SD), and the data were considered statistically significant when p < 0.05. Statistical significance is represented in figures by * p < 0.05, ** p < 0.005, *** p < 0.0005 or **** p < 0.0005.

## 3. Results

### 3.1. Lurasidone and olanzapine exert contrasting effects on gut microbiota composition and diversity

The dosage regimen for lurasidone and olanzapine (*i.e.,* 7.5 mg/kg for 21 days) were selected based on prior literature, aiming to achieve optimal D2 receptor occupancy (75%) and thus emulate normal physiological response (Mann et al., 2013; Qian et al., 2023). Stool samples were gathered on Day 0 (prior to the initial dose), weekly thereafter, and on Day 21 (following the last dose), with changes in relative microbial abundance at the family taxonomic level for each animal highlighted in **Fig. 3A-B** with grouped data on relative abundance also noted in **Fig. 3C-D**. Notably, *Lachnospiraceae* and *Bacteroidaceae* decreased in relative abundance following olanzapine dosing. Both families are widely involved in pathways of SCFA production and harbour functional activity genes which contribute to glucose and starch degradation (Biddle et al., 2013). The observed increase in relative abundance of *Lactobacillaceae* and *Peptostreptococcaceae* in groups exposed to olanzapine may be opportunistic. Lurasidone exerted positive changes to the gut microbiota at the family taxa level, with increases in *Lachnospiraceae* relative abundance and maintenance of the baseline healthy ecosystem.

Observed changes in alpha diversity (Shannon’s Index) and total operational taxonomical units (OTUs) are presented in **Fig. 2A-C** and **Fig. 2B-D** for lurasidone and olanzapine treated groups, respectively. Shannon’s index analysis illustrates both abundance and uniformity of species distribution, with results indicating lurasidone as potentiating a more diverse gut microbiota ecosystem (+0.5 increase in Shannon’s index) compared to olanzapine (−1 decrease). Lurasidone exerted positive impacts on the gut microbiota, through enhancement of OTUs (+50 OTUs) suggesting increased diversity. Olanzapine exhibited the contrasting effect with mean decrease of –75 OTUs suggesting greater dominance by fewer taxa. Noteworthy is the rapidity and linearity of changes in OTUs and Shannon’s index for both lurasidone and olanzapine, with the observed alterations occurring within 7 days and escalating throughout the 21-day period. Likewise, principal component analysis (PCoA) revealed a significant and contrasting shift between day 0 and day 21 in the gut microbial compositions for animals administered with olanzapine, as highlighted by separation of microbial components through beta diversity (**Fig. 2F**).

**Figure 1:**
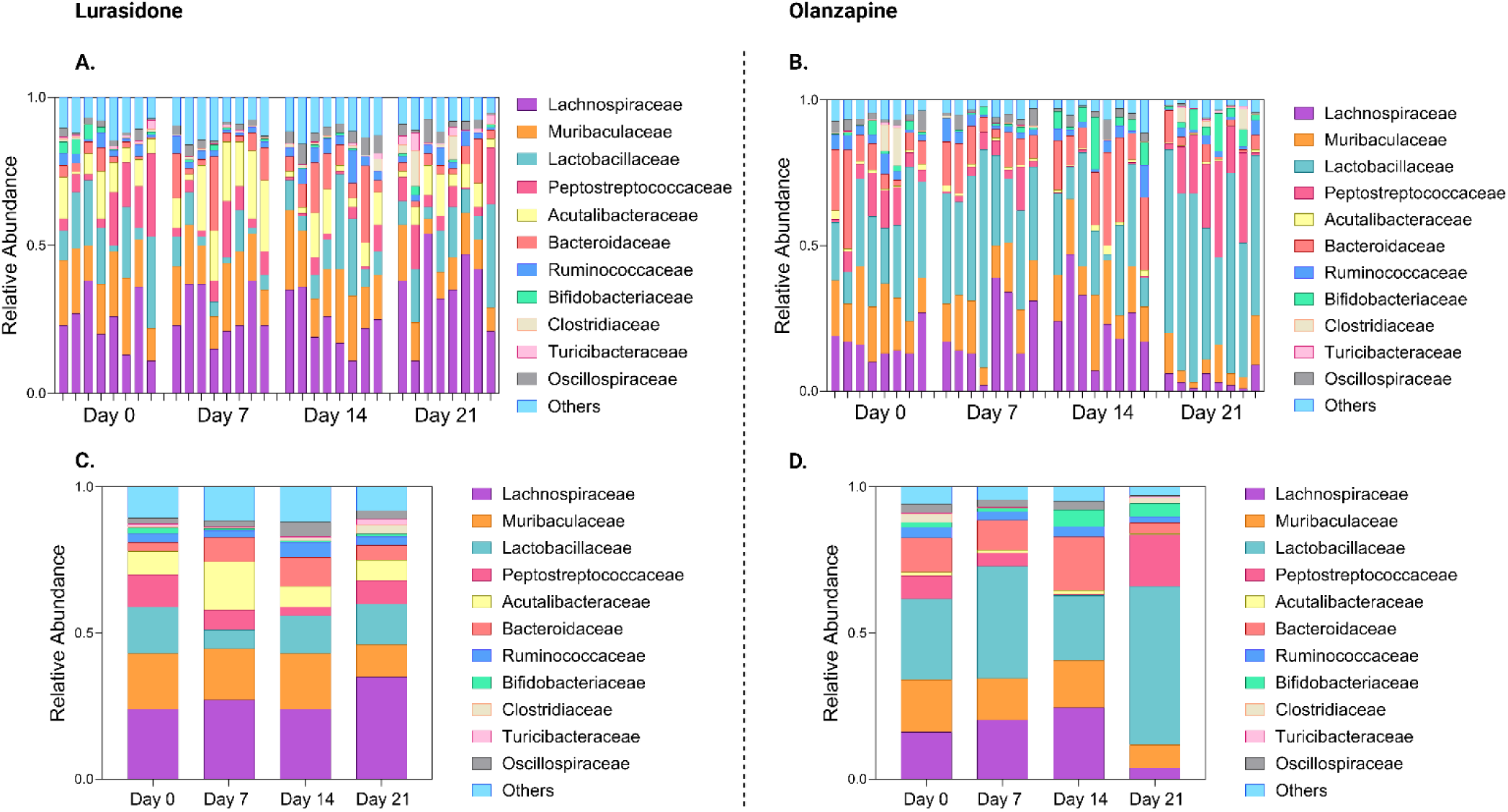
Panels **A.** and **B.** showcase changes to relative abundance of gut microbiota families post exposure to lurasidone and olanzapine, respectively. Panels **C.** and **D.** highlight grouped data on relative abundance for lurasidone and olanzapine respectively.

**Figure 2:**
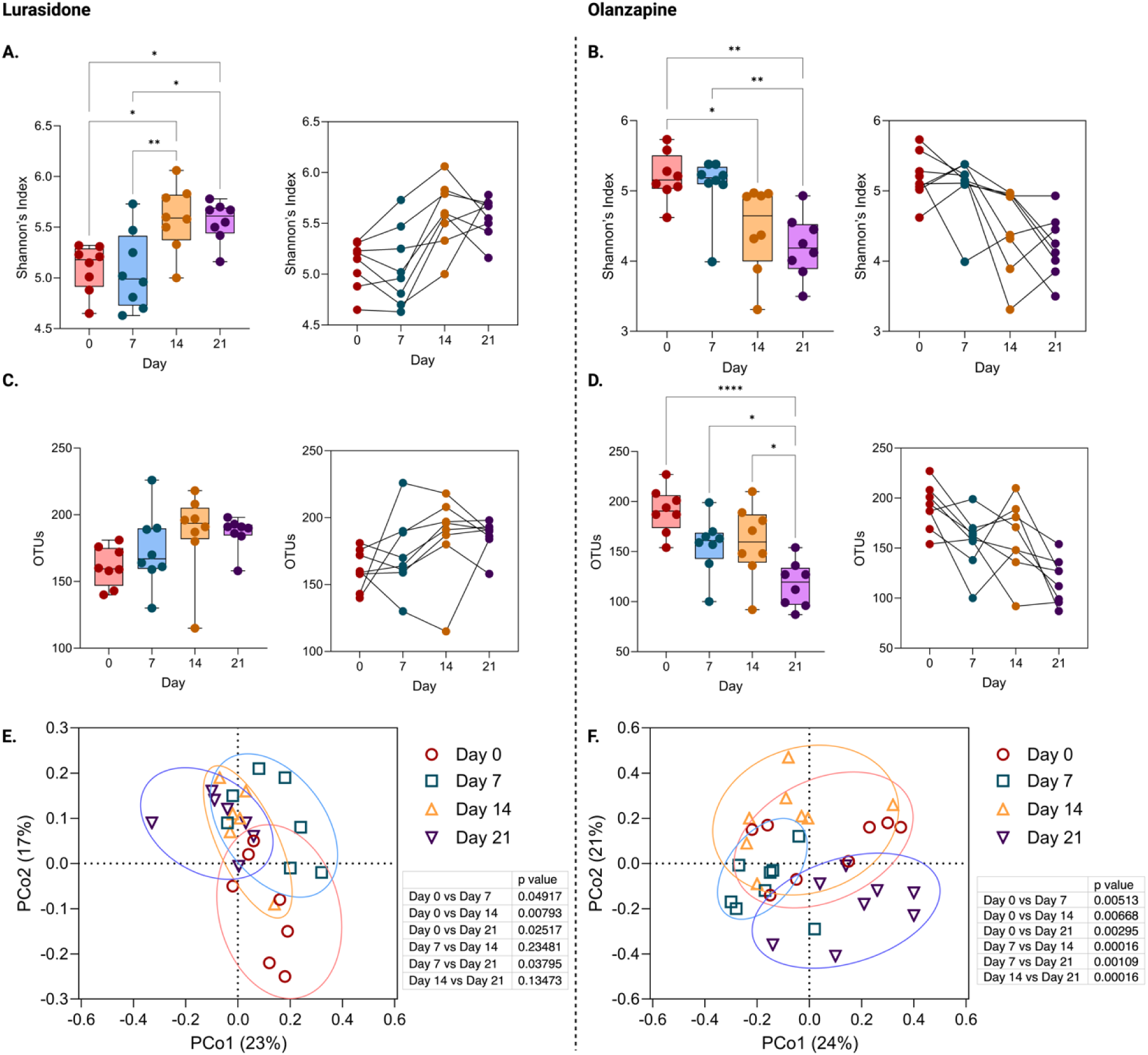
Lurasidone and Olanzapine triggered contrasting impacts on the gut microbiota with olanzapine decreasing microbiota diversity whilst lurasidone largely maintains a positive profile. Panels **A.** and **B.** showcase the contrasting impacts of olanzapine and lurasidone on Shannon’s index with olanzapine significantly and consistently decreasing gut microbiota ‘richness’ whilst lurasidone improves gut microbiota diversity. Panels **C.** and **D.** showcase decreases in total OTUs for olanzapine whilst lurasidone increased total OTUs. Panels **E.** and **F.** showcase shifts in microbiota composition through principal component analyses for lurasidone and olanzapine respectively

Alongside adverse alterations to alpha and beta diversity, olanzapine also triggered a rise in the *Firmicutes*: *Bacteroidetes* (F: B) ratio (5-fold increase) (**Fig. 3A**), a potent indicator of gut microbiome dysbiosis. Lurasidone did not significant alter F: B ratios from baseline levels. Notable is the abruptness in increase, with majority of the changes occurring over the last 7-day period of the trial. This sudden rise coincidences with the cumulating decreases in key taxa negatively correlated with weight gain, namely *Lachnoclostridium*, *Parasutterella* and *Blautia* (all of which experienced approximate halving in populations) (**Fig. C-H** respectively) suggesting a threshold level had been reached. In contrast, lurasidone did not trigger any statistically significant changes to both F: B ratio nor the aforementioned key taxa.

**Figure 3:**
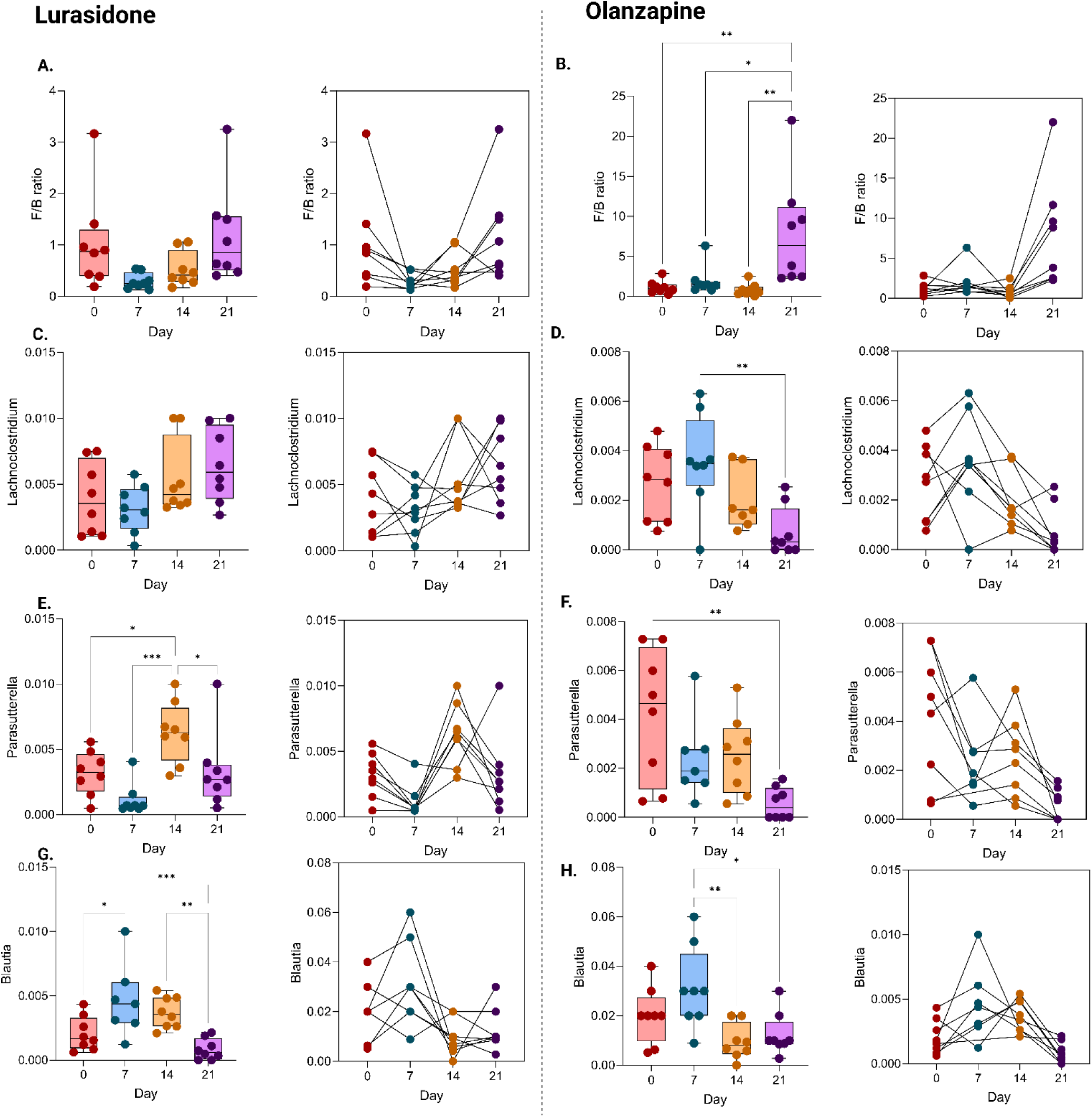
Panels **A.** and **B.** showcase changes to the Firmicutes: Bacteroidetes (F: B) ratio upon exposure to lurasidone and olanzapine respectively. Olanzapine triggers a statistically significant increase in F: B ratio, particularly on day 21. Comparisons of lurasidone and olanzapine reveal contrasting impacts on key taxa including Lachnoclostridium (panels **C.** and **D.**) Parasutterella (panels **E.** and **F.**) and Blautia (panels **G.** and **H.**) with olanzapine consistently decreasing relative abundance.

### 3.2. Lurasidone does not induce metabolic dysfunction through weight gain, inflammation or changes to blood triglycerides and glycaemic levels

Lurasidone showcased minimal impact on metabolic markers measured (no statistically significant impacts on weight gain or inflammatory and metabolic markers) (**Fig. 4B-D**). In contrast, olanzapine triggered metabolic dysfunction as highlighted by statistically significant increases in TNF-α concentrations (2-fold increase), blood triglycerides (2-fold increase) and blood glucose (3-fold increase) (**Fig. 4B, E, F**). Comparable to the abrupt olanzapine-induced increase in F: B ratio observed in **Fig. 3B**, blood triglycerides and blood glucose levels similarly follow the same abrupt increase in the last 7 days of the trial, with significant hyperglycaemia observed after 21 days (**Fig. 4E**). These alterations cumulate to potentiate weight gain, as observed in **Fig. 4A**, with olanzapine showcasing a statistically significant increase in weight (+30%) compared to lurasidone and control (+10%).

**Figure 4:**
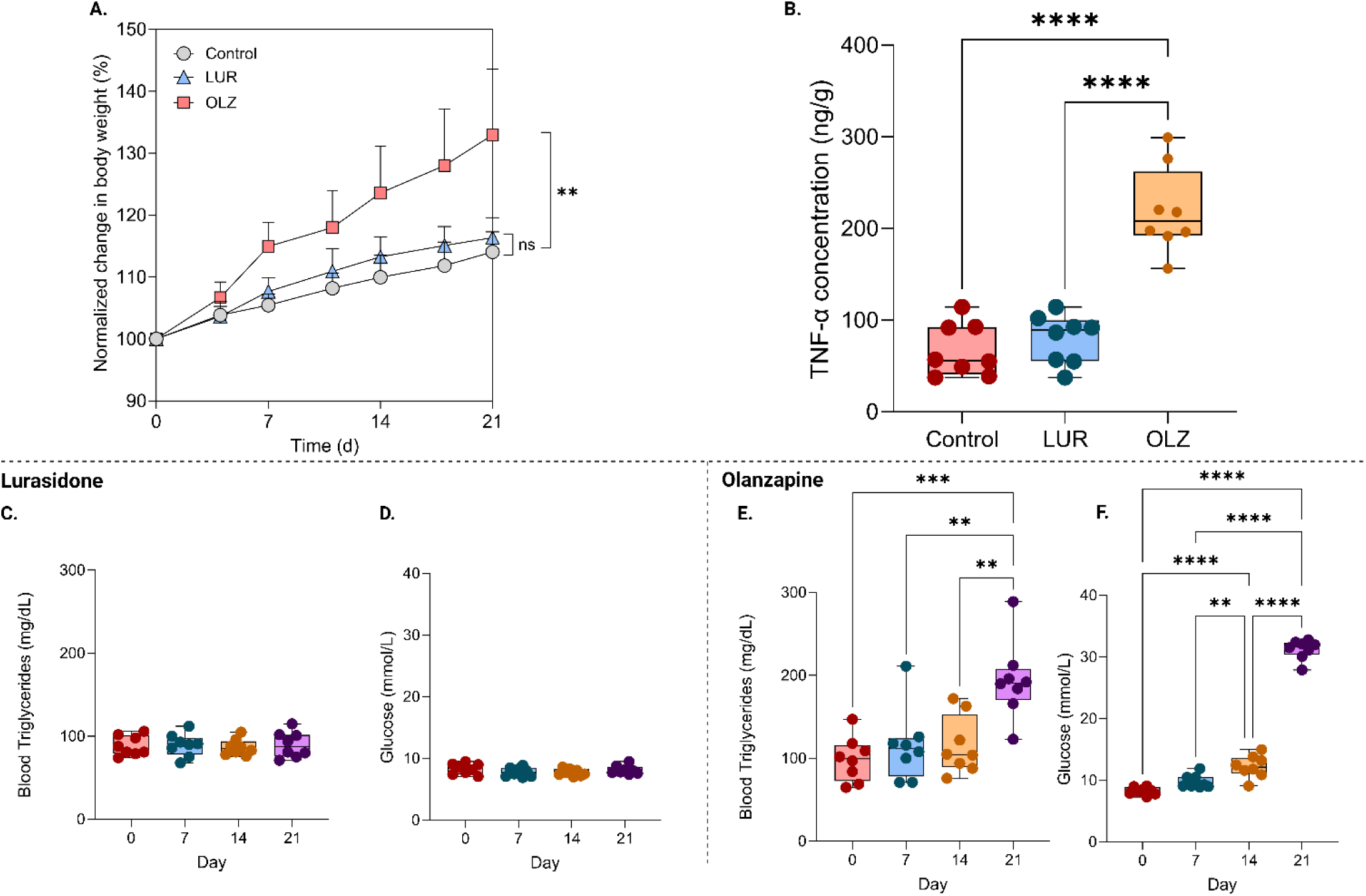
**A.** Lurasidone showed a non-significant increase in body weight over time compared to control however olanzapine exposed rats experienced a 30% increase in body weight. **B.** TNF-α concentrations showcased a statistically significant rise in presence of olanzapine. Panels **C** and **D** highlight impact the lack of impacts of lurasidone on blood triglycerides and blood glucose levels respectively. Olanzapine did create statistically significant impacts, as highlighted in panels **E.** and **F.** whereby both blood triglycerides and blood glucose experienced substantial increases.

### 3.3. Olanzapine-induced metabolic dysfunction and intestinal inflammation correlates with gut dysbiosis

As depicted in **Fig. 1** and **Fig. 3**, olanzapine induced dysbiosis across several families of gut microbiota. Further, the abundance of key genera, including *Lachnoclostridium*, *Parasutterella* and *Blautia* were notably affected, with all showing statistically significant decreases post olanzapine exposure for 21 days. The abundance of all three genera negatively correlated with body weight increase (R^2^ = 0.44, 0.46 and 0.33, respectively). The importance of the F: B ratio is highlighted in **Fig. 5A-C**, whereby olanzapine-induced changes to the F: B ratio correlated strongly with body weight increase (R^2^ = 0.76), and also showcased trends with blood glucose (R^2^ = 0.43) and blood triglyceride levels (R^2^ = 0.11). Lurasidone and its neutrality towards the F: B ratio did not significantly impact any of the aforementioned factors.

**Figure 5:**
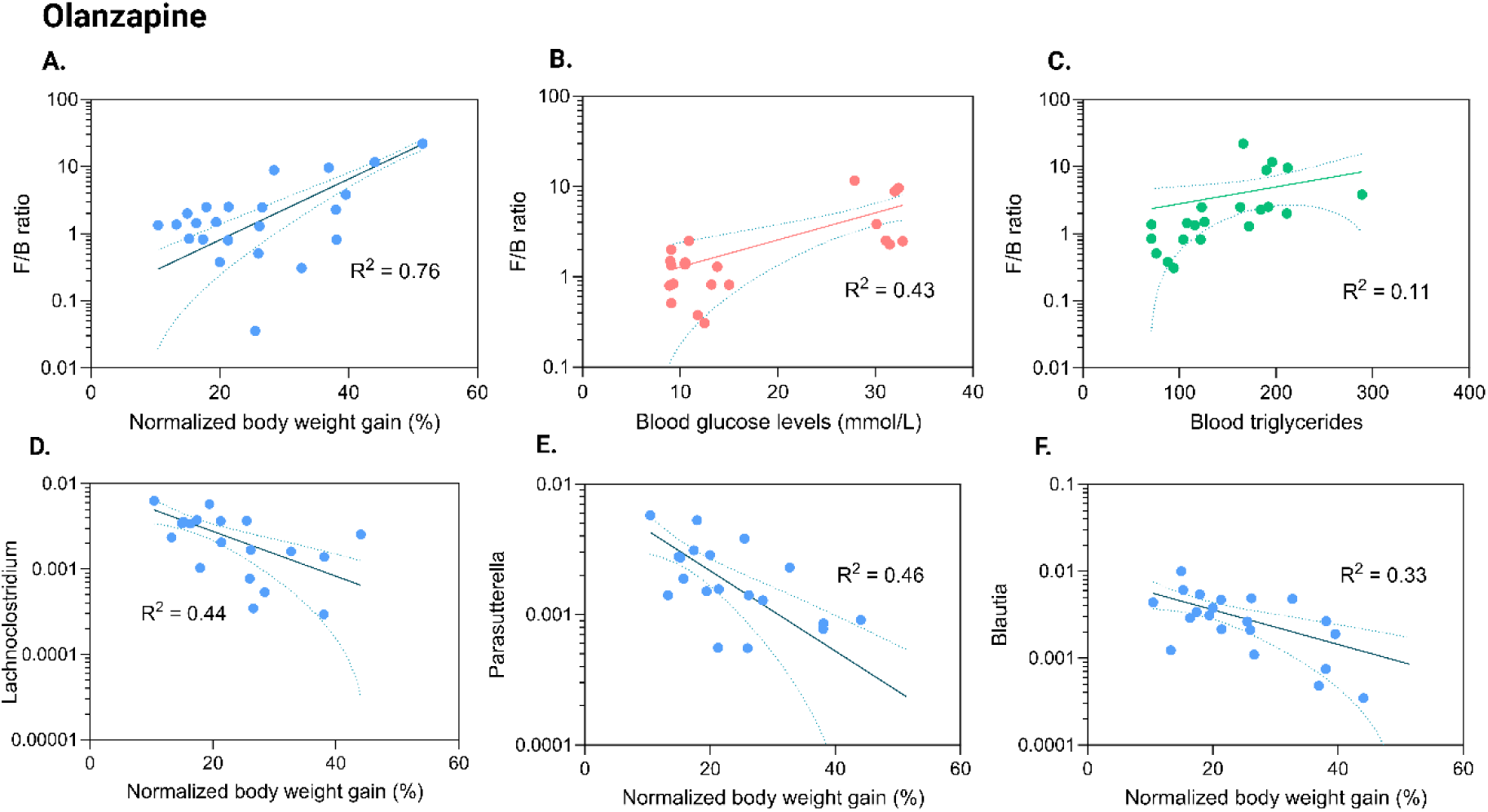
Panels **A-C.** explore the varying correlations between F: B ratios in olanzapine treated rodents and observed impacts on body weight, blood glucose and blood triglyceride levels respectively. Panels **D-F.** highlight various families impacted by olanzapine administration (Lachnoclostridium, Parasutterella and Blautia respectively) and their correlation with body weight gain.

## 4. Discussion

There exist concerns over antipsychotic-induced gut microbiota dysbiosis leading to metabolic dysfunction, weight gain and impairing therapeutic outcomes. Commonly prescribed atypical antipsychotics such as olanzapine are often associated with rapid and pronounced metabolic dysfunction, with clinical data suggesting body weight to increase by +0.9 kg per month up to +6 to +10 kg after 1 year of treatment (Nemeroff, 1997). The underlying mechanism behind this is suggested to be due to changes to the gut microbiota which potentiate an obesogenic profile. This has led to the proposition that all antipsychotics are detrimental to metabolic functioning and the gut microbiota (Davey et al., 2013; Hao et al., 2023; Minichino et al., 2023; Munawar et al., 2021; Pochiero et al., 2021; Qian et al., 2023; Seeman, 2021). Lurasidone, another atypical second-generation antipsychotic, has received lesser investigation with its impact of body weight and metabolic function not fully understood. Findings from studies exploring the impact of lurasidone on weight gain have varied, from weight increases, neutrality and decreases all being observed (Dayabandara et al., 2017; Greger et al., 2021; Loebel et al., 2014; Meyer et al., 2015, 2017; Ng-Mak et al., 2018; Pochiero et al., 2021; Reynolds et al., 2019). This discrepancy has led to inconclusive opinions on lurasidone weight gain, rather being referred to as a ‘less potentiating’ agent of weight gain compared to alternative antipsychotics (Greger et al., 2021; Meyer et al., 2017; Pochiero et al., 2021).

This novel study challenges and dispels the notion that all antipsychotics negatively disrupt the gut microbiota and explains a rationale for inconsistency in findings for lurasidone-induced weight gain. Through contrasting the impacts of lurasidone and olanzapine on the gut microbiota, the present study suggests that the gut microbiota might have a significant impact on antipsychotic-induced obesity, specifically explaining lurasidone’s weight and metabolic neutrality through its observed positive impacts on the gut microbiota.

Olanzapine outcomes in this study are consistent with prior work with similarly significant increase in weight and disturbance in metabolic health and gut microbiota compositions (Cussotto et al., 2021, 2021; Davey et al., 2013; A. C.-C. Kao, Spitzer, et al., 2018; Leucht et al., 2013; Qian et al., 2023; Ratzoni et al., 2002). Rats exposed to olanzapine showcased a 30% increase in body weight compared to control which is marginally higher than that observed in studies by Qian et al. (Qian et al., 2023). Noteworthy is the rapidity and linearity in rate of olanzapine-induced weight gain with consistent weight gain over the observed 21 days. This corroboration extends to similarities in gut compositional changes with olanzapine triggering dynamic shifts in gut microbial profiles, resulting in decreases to Shannon’s index and beta diversity comparable to that observed by Qian et al. (Qian et al., 2023).

Bacterial families key to metabolic health were inhibited by olanzapine, including *Lachnospiraceae* and *Bacteroidaceae*. *Lachnospiraceae* and its associated repertoire of enzymes are implicated in the pathways for SCFA production (propionate, butyrate, and acetate), alongside metabolism of carbohydrates (namely L-fucose, starch, insulin and arabinoxylan) and aromatic amino acids (resulting in production of indole, 3-indolepropionic acid, para-cresol, and phenol) (Vacca et al., 2020). *Bacteroidaceae* harbour polysaccharide utilisation loci (PULs) necessary to provide protein machinery for metabolism of carbohydrates (Zafar & Saier, 2021). One such PUL is the starch utilisation system which includes numerous cell-surface proteins and enzymes necessary for breakdown for starch and related-products (Tuson et al., 2018). This PUL machinery also allows inter-species cross-feeding, for example provision of starch metabolism-related products such as glucose and maltose to neighbouring *Eubacterium ramulus* which in turn metabolise flavonoids such as quercetin for use by *Bacteroidaceae* (Zafar & Saier, 2021). Administration of olanzapine led to significant decreases relative abundances for both populations, which is expected to result in decreased capacity for SCFA production, improper metabolism of ingested nutrients and damage to ecosystem-level commensal nutrient exchange (Cussotto et al., 2019). This is suggested to be the basis for its observed impact on weight gain and metabolic health and corroborated by Davey et al. (Davey et al., 2013). Furthermore, our findings suggest an increase in relative abundance of *Lactobacillaceae* however, despite its non-pathogenic background it has been reported to be opportunistic by nature (Batt & Tortorello, 2014). Thus, the increased relative abundance appears to be an adaptation by *Lactobacillaceae* and its surrounding ecosystem rather than directly mediated by olanzapine.

Similar exploration of the impact of olanzapine and lurasidone on *Lachnoclostridium*, *Parasutterella* and *Blautia* genera illustrates an underlying mechanism for observed weight gain discrepancies. Specifically, decrease in relative abundance of three families showcased casual trends with weight gain (R^2^ = 0.44, 0.46 and 0.33 respectively), and all were significantly inhibited by olanzapine (−50%, −60% and −50% decrease in relatively abundance respectively). Lurasidone did not significantly impact the relative abundance of these noteworthy taxa, and contrastingly to olanzapine, positively increased *Lachnoclostridium* abundance, which may garner protective properties against weight gain. Furthermore, olanzapine also altered *Firmicutes*: *Bacteroidetes* (F: B) ratio, a commonly reported signature of gut microbiota dysbiosis (Koliada et al., 2017; Magne et al., 2020). This ratio correlated strongly (R^2^ = 0.76) with observed weight gain. Similarly, the olanzapine-altered F: B ratio (5-fold increase) also showed causal trends with metabolic markers measured, namely blood triglycerides and blood glucose levels (R^2^ = 0.43 0.11 respectively). When considered with other olanzapine-induced changes it may suggest the underpinning mechanism behind the observed 2-fold increase in blood triglycerides and 3-fold increase in blood glucose levels alongside 30% increase in weight gain experienced by olanzapine treatment groups but not by lurasidone groups.

Novel in this study is the exploration of lurasidone and its impact on gut and metabolic health. Lurasidone showcased non-significant weight gain compared to control groups, suggesting it to be weight neutral. This statistically non-significant result aligns with prior literature by Greger et al. which reported a statically non-significant mean −0.27 kg weight loss after treatment with lurasidone for one year (Greger et al., 2021). This theme of neutrality extends across all measured parameters. The gut microbiota displayed largely positive to neutral impacts with dynamic improvements in OTUs and microbial richness and diversity, suggesting lurasidone had a positive bearing on gut functioning. These novel findings suggest lurasidone to be at least ‘gut neutral’ if not beneficial for the gut microbiome. Furthermore, contrastive to olanzapine, F: B ratio was not significantly impacted nor the relative abundance with any key families associated with weight gaining propensities. Instead, lurasidone positively modulated *Lachnospiraceae* relative abundance in stark contrast to olanzapine which is suggestive of positive gut functioning.

Similarly, lurasidone-induced impacts on all metabolic markers measured including TNF-α concentrations, blood total triglycerides and blood glucose, were negligible and statistically non-significant. These novel findings suggest lurasidone to be metabolically neutral, which when compared to olanzapine groups propose gut-positive impacts to be the underpinning mechanism.

This finding has potentially significant impacts not just in promoting the favourability of lurasidone in weight or metabolically sensitive patients (for example those with comorbidities such as diabetes mellitus), but also paves a plausible underlying mechanism for weight neutrality observed in other atypical antipsychotics such as ziprasidone (Park et al., 2013). Given dysbiosis is implicated with a multitude of various adverse health impacts ranging from metabolic dysfunction to worsening of mental illness, lurasidone’s relative gut neutrality is suggested to be favourable (Magne et al., 2020; Munawar et al., 2021). Furthermore, it allows for unhindered support from additional gut-targeted interventions such as pro-, pre-, syn- and post-biotics to be utilised to further guide the gut microbiota and non-pharmacologically benefit the patient (Burnet, 2015; Kamath et al., 2023; A. C. C. Kao et al., 2016; A. C.-C. Kao, Burnet, et al., 2018; Radford-Smith & Anthony, 2023). Following this, the present investigation lays the groundwork for clinical trials to determine whether the findings observed in rodents correlate with the effects of lurasidone on the human gut microbiome. Further work is required to investigate if lurasidone’s positive impacts on the gut microbiota are direct or indirectly achieved through alterations higher up in the hierarchy and then transmitted through the gut-brain axis.

Finally, the study highlights need for further exploration into the varying impacts of antipsychotics on the gut and metabolic health, dissuading the notion that most antipsychotics uniformly and adversely disrupt the gut microbiota (Cussotto et al., 2019; Davey et al., 2013; Greger et al., 2021; Hao et al., 2023; A. C.-C. Kao, Spitzer, et al., 2018, 2018; Minichino et al., 2023; Munawar et al., 2021). With such dissimilarity in impacts, it may necessitate re-evaluation of treatment regimens for schizophrenia to accommodate for sensitivities, such as to metabolic fluctuations. Given there is mounting evidence for the metabolically disrupting tendencies of certain antipsychotics such as olanzapine, it is anticipated that future studies uncover better-suited alternatives such as lurasidone and allow for personalised medicines based on propensity for weight gain and comorbidities. It is also anticipated that such encouraging results for lurasidone may drive necessary work required promote its place in standard therapy.

## 5. Conclusions

This novel study explores the complex interplay between antipsychotic medications, gut microbiota dynamics, and metabolic health. By contrasting the effects of olanzapine and lurasidone on weight gain and gut microbiota composition, it challenges the prevailing notion that all antipsychotics uniformly disrupt metabolic health. While olanzapine demonstrates significant weight gain and dysbiosis, lurasidone does not exhibit significant impacts on weight and metabolic parameters measured, suggesting a unique “gut neutrality” property. These findings not only advocate for the preferential use of lurasidone, especially in weight or metabolically sensitive patients, but also provide a rationale for exploring similar effects in other antipsychotics like ziprasidone. Furthermore, the study underscores the importance of a personalised medicines approach in schizophrenia (and more broadly, mental health) treatment regimens to mitigate metabolic disturbance, with special awareness of gut-targeted therapies. This research lays the groundwork for future clinical trials to validate these findings in human subjects and emphasises imperative for further exploration into the varying impacts of antipsychotics on the gut microbiota.

## Acknowledgements

The Hospital Research Foundation (THRF) Group are gratefully acknowledged for their EMCR Fellowship funding and generous support provided to Dr Paul Joyce (2022-CF-EMCR-004-25314). The research team would like to kindly acknowledge Dr. Aurelia Elz for her expertise and assistance in undertaking in vivo evaluations in this study.

## References

Ågren, R. (2021). Worldwide antipsychotic drug search intensities: Pharmacoepidemological estimations based on Google Trends data. Scientific Reports, 11(1), 13136. 10.1038/s41598-021-92204-0

Ait Chait, Y., Mottawea, W., Tompkins, T. A., & Hammami, R. (2021). Nutritional and therapeutic approaches for protecting human gut microbiota from psychotropic treatments. Progress in Neuro-Psychopharmacology and Biological Psychiatry, 108, 110182. 10.1016/j.pnpbp.2020.110182

Ait Chait, Y., Mottawea, W., Tompkins, T. A., & Hammami, R. (2023). Evidence of the Dysbiotic Effect of Psychotropics on Gut Microbiota and Capacity of Probiotics to Alleviate Related Dysbiosis in a Model of the Human Colon. International Journal of Molecular Sciences, 24(8), 7326. 10.3390/ijms24087326

Aoun, A., Darwish, F., & Hamod, N. (2020). The Influence of the Gut Microbiome on Obesity in Adults and the Role of Probiotics, Prebiotics, and Synbiotics for Weight Loss. Preventive Nutrition and Food Science, 25(2), 113–123. 10.3746/pnf.2020.25.2.113

Appleton, J. (2018). The Gut-Brain Axis: Influence of Microbiota on Mood and Mental Health. Integrative Medicine: A Clinician’s Journal, 17(4), 28.

Batt, C. A., & Tortorello, M. L. (Eds.). (2014). Encyclopedia of food microbiology (2. ed). AP, Academic Press/Elsevier.

Bear, T., Dalziel, J., Coad, J., Roy, N., Butts, C., & Gopal, P. (2021). The Microbiome-Gut-Brain Axis and Resilience to Developing Anxiety or Depression under Stress. Microorganisms, 9(4), Article 4. 10.3390/microorganisms9040723

Biddle, A., Stewart, L., Blanchard, J., & Leschine, S. (2013). Untangling the Genetic Basis of Fibrolytic Specialization by Lachnospiraceae and Ruminococcaceae in Diverse Gut Communities. Diversity, 5(3), 627–640. 10.3390/d5030627

Burnet, P. W. (2015). S.20.02 Prebiotics and anxiety: Translational studies toward new therapeutic strategies. European Neuropsychopharmacology, 25, S141. 10.1016/S0924-977X(15)30097-3

Clapp, M., Aurora, N., Herrera, L., Bhatia, M., Wilen, E., & Wakefield, S. (2017). Gut Microbiota’s Effect on Mental Health: The Gut-Brain Axis. Clinics and Practice, 7(4), 987. 10.4081/cp.2017.987

Cussotto, S., Clarke, G., Dinan, T. G., & Cryan, J. F. (2019). Psychotropics and the Microbiome: A Chamber of Secrets…. Psychopharmacology, 236(5), 1411–1432. 10.1007/s00213-019-5185-8

Cussotto, S., Walsh, J., Golubeva, A. V., Zhdanov, A. V., Strain, C. R., Fouhy, F., Stanton, C., Dinan, T. G., Hyland, N. P., Clarke, G., Cryan, J. F., & Griffin, B. T. (2021). The gut microbiome influences the bioavailability of olanzapine in rats. EBioMedicine, 66, 103307. 10.1016/j.ebiom.2021.103307

Davey, K. J., Cotter, P. D., O’Sullivan, O., Crispie, F., Dinan, T. G., Cryan, J. F., & O’Mahony, S. M. (2013). Antipsychotics and the gut microbiome: Olanzapine-induced metabolic dysfunction is attenuated by antibiotic administration in the rat. Translational Psychiatry, 3(10), e309–e309. 10.1038/tp.2013.83

Dayabandara, M., Hanwella, R., Ratnatunga, S., Seneviratne, S., Suraweera, C., & De Silva, V. (2017). Antipsychotic-associated weight gain: Management strategies and impact on treatment adherence. *Neuropsychiatric Disease and Treatment*, Volume 13, 2231– 2241. 10.2147/NDT.S113099

Greger, J., Aladeen, T., Lewandowski, E., Wojcik, R., Westphal, E., Rainka, M., & Capote, H. (2021). Comparison of the Metabolic Characteristics of Newer Second Generation Antipsychotics: Brexpiprazole, Lurasidone, Asenapine, Cariprazine, and Iloperidone With Olanzapine as a Comparator. Journal of Clinical Psychopharmacology, 41(1), 5–12. 10.1097/JCP.0000000000001318

Hao, S., Zhou, Y., Zhang, X., & Jiang, H. (2023). Gut microbiome profiles may be related to atypical antipsychotic associated overweight in Asian children with psychiatric disorder: A preliminary study. Frontiers in Cellular and Infection Microbiology, 13, 1124846. 10.3389/fcimb.2023.1124846

Kamath, S., Stringer, A. M., Prestidge, C. A., & Joyce, P. (2023). Targeting the gut microbiome to control drug pharmacomicrobiomics: The next frontier in oral drug delivery. Expert Opinion on Drug Delivery, 1–17. 10.1080/17425247.2023.2233900

Kao, A. C. C., Harty, S., & Burnet, P. W. J. (2016). The Influence of Prebiotics on Neurobiology and Behavior. In International Review of Neurobiology (Vol. 131, pp. 21–48). Elsevier. 10.1016/bs.irn.2016.08.007

Kao, A. C.-C., Burnet, P. W. J., & Lennox, B. R. (2018). Can prebiotics assist in the management of cognition and weight gain in schizophrenia? Psychoneuroendocrinology, 95, 179–185. 10.1016/j.psyneuen.2018.05.027

Kao, A. C.-C., Spitzer, S., Anthony, D. C., Lennox, B., & Burnet, P. W. J. (2018). Prebiotic attenuation of olanzapine-induced weight gain in rats: Analysis of central and peripheral biomarkers and gut microbiota. Translational Psychiatry, 8(1), 66. 10.1038/s41398-018-0116-8

Koliada, A., Syzenko, G., Moseiko, V., Budovska, L., Puchkov, K., Perederiy, V., Gavalko, Y., Dorofeyev, A., Romanenko, M., Tkach, S., Sineok, L., Lushchak, O., & Vaiserman, A. (2017). Association between body mass index and Firmicutes/Bacteroidetes ratio in an adult Ukrainian population. BMC Microbiology, 17(1), 120. 10.1186/s12866-017-1027-1

Leucht, S., Cipriani, A., Spineli, L., Mavridis, D., Örey, D., Richter, F., Samara, M., Barbui, C., Engel, R. R., Geddes, J. R., Kissling, W., Stapf, M. P., Lässig, B., Salanti, G., & Davis, J. M. (2013). Comparative efficacy and tolerability of 15 antipsychotic drugs in schizophrenia: A multiple-treatments meta-analysis. The Lancet, 382(9896), 951– 962. 10.1016/S0140-6736(13)60733-3

Loebel, A., Cucchiaro, J., Silva, R., Kroger, H., Hsu, J., Sarma, K., & Sachs, G. (2014). Lurasidone Monotherapy in the Treatment of Bipolar I Depression: A Randomized, Double-Blind, Placebo-Controlled Study. American Journal of Psychiatry, 171(2), 160–168. 10.1176/appi.ajp.2013.13070984

Magne, F., Gotteland, M., Gauthier, L., Zazueta, A., Pesoa, S., Navarrete, P., & Balamurugan, R. (2020). The Firmicutes/Bacteroidetes Ratio: A Relevant Marker of Gut Dysbiosis in Obese Patients? Nutrients, 12(5), 1474. 10.3390/nu12051474

Mann, S., Chintoh, A., Giacca, A., Fletcher, P., Nobrega, J., Hahn, M., & Remington, G. (2013). Chronic olanzapine administration in rats: Effect of route of administration on weight, food intake and body composition. Pharmacology Biochemistry and Behavior, 103(4), 717–722. 10.1016/j.pbb.2012.12.002

Meltzer, H. Y., Cucchiaro, J., Silva, R., Ogasa, M., Phillips, D., Xu, J., Kalali, A. H., Schweizer, E., Pikalov, A., & Loebel, A. (2011). Lurasidone in the Treatment of Schizophrenia: A Randomized, Double-Blind, Placebo- and Olanzapine-Controlled Study. American Journal of Psychiatry, 168(9), 957–967. 10.1176/appi.ajp.2011.10060907

Meyer, J. M., Mao, Y., Pikalov, A., Cucchiaro, J., & Loebel, A. (2015). Weight change during long-term treatment with lurasidone: Pooled analysis of studies in patients with schizophrenia. International Clinical Psychopharmacology, 30(6), 342–350. 10.1097/YIC.0000000000000091

Meyer, J. M., Ng-Mak, D. S., Chuang, C.-C., Rajagopalan, K., & Loebel, A. (2017). Weight changes before and after lurasidone treatment: A real-world analysis using electronic health records. Annals of General Psychiatry, 16(1), 36. 10.1186/s12991-017-0159-x

Minichino, A., Preston, T., Fanshawe, J. B., Fusar-Poli, P., McGuire, P., Burnet, P. W. J., & Lennox, B. R. (2023). Psycho-pharmacomicrobiomics: A Systematic Review and Meta-analysis. Biological Psychiatry, S0006322323014865. 10.1016/j.biopsych.2023.07.019

Morgan, A. P., Crowley, J. J., Nonneman, R. J., Quackenbush, C. R., Miller, C. N., Ryan, A. K., Bogue, M. A., Paredes, S. H., Yourstone, S., Carroll, I. M., Kawula, T. H., Bower, M. A., Sartor, R. B., & Sullivan, P. F. (2014). The Antipsychotic Olanzapine Interacts with the Gut Microbiome to Cause Weight Gain in Mouse. PLoS ONE, 9(12), e115225. 10.1371/journal.pone.0115225

Munawar, N., Ahsan, K., Muhammad, K., Ahmad, A., Anwar, M. A., Shah, I., Al Ameri, A. K., & Al Mughairbi, F. (2021). Hidden Role of Gut Microbiome Dysbiosis in Schizophrenia: Antipsychotics or Psychobiotics as Therapeutics? International Journal of Molecular Sciences, 22(14), 7671. 10.3390/ijms22147671

Nemeroff, C. B. (1997). Dosing the antipsychotic medication olanzapine. The Journal of Clinical Psychiatry, 58 *Suppl 10*, 45–49.

Ng-Mak, D., Tongbram, V., Ndirangu, K., Rajagopalan, K., & Loebel, A. (2018). Efficacy and metabolic effects of lurasidone versus brexpiprazole in schizophrenia: A network meta-analysis. Journal of Comparative Effectiveness Research, 7(8), 737–748. 10.2217/cer-2018-0016

Park, S., Yi, K. K., Kim, M.-S., & Hong, J. P. (2013). Effects of ziprasidone and olanzapine on body composition and metabolic parameters: An open-label comparative pilot study. Behavioral and Brain Functions, 9(1), 27. 10.1186/1744-9081-9-27

Pochiero, I., Calisti, F., Comandini, A., Del Vecchio, A., Costamagna, I., Rosignoli, M. T., Cattaneo, A., Nunna, S., Peduto, I., Heiman, F., Chang, H.-C., Chen, C.-C., & Correll, C. (2021). Impact of Lurasidone and Other Antipsychotics on Body Weight: Real-World, Retrospective, Comparative Study of 15,323 Adults with Schizophrenia. International Journal of General Medicine, Volume 14, 4081–4094. 10.2147/IJGM.S320611

Qian, L., He, X., Liu, Y., Gao, F., Lu, W., Fan, Y., Gao, Y., Wang, W., Zhu, F., Wang, Y., & Ma, X. (2023). Longitudinal Gut Microbiota Dysbiosis Underlies Olanzapine-Induced Weight Gain. Microbiology Spectrum, 11(4), e00058–23. 10.1128/spectrum.00058-23

Radford-Smith, D. E., & Anthony, D. C. (2023). Prebiotic and Probiotic Modulation of the Microbiota–Gut–Brain Axis in Depression. Nutrients, 15(8), 1880. 10.3390/nu15081880

Radford-Smith, D. E., Probert, F., Burnet, P. W. J., & Anthony, D. C. (2022). Modifying the maternal microbiota alters the gut–brain metabolome and prevents emotional dysfunction in the adult offspring of obese dams. Proceedings of the National Academy of Sciences, 119(9), e2108581119. 10.1073/pnas.2108581119

Ratzoni, G., Gothelf, D., Brand-Gothelf, A., Reidman, J., Kikinzon, L., Gal, G., Phillip, M., Apter, A., & Weizman, R. (2002). Weight Gain Associated With Olanzapine and Risperidone in Adolescent Patients: A Comparative Prospective Study. Journal of the American Academy of Child & Adolescent Psychiatry, 41(3), 337–343. 10.1097/00004583-200203000-00014

Reynolds, G. P., Dalton, C. F., Watrimez, W., Jackson, J., & Harte, M. K. (2019). Adjunctive Lurasidone Suppresses Food Intake and Weight Gain Associated with Olanzapine Administration in Rats. Clinical Psychopharmacology and Neuroscience, 17(2), 314–317. 10.9758/cpn.2019.17.2.314

Safadi, J. M., Quinton, A. M. G., Lennox, B. R., Burnet, P. W. J., & Minichino, A. (2022). Gut dysbiosis in severe mental illness and chronic fatigue: A novel trans-diagnostic construct? A systematic review and meta-analysis. Molecular Psychiatry, 27(1), 141–153. 10.1038/s41380-021-01032-1

Sarkar, A., Harty, S., Johnson, K. V.-A., Moeller, A. H., Carmody, R. N., Lehto, S. M., Erdman, S. E., Dunbar, R. I. M., & Burnet, P. W. J. (2020). The role of the microbiome in the neurobiology of social behaviour. Biological Reviews, 95(5), 1131–1166. 10.1111/brv.12603

Sarkar, A., Lehto, S. M., Harty, S., Dinan, T. G., Cryan, J. F., & Burnet, P. W. J. (2016). Psychobiotics and the Manipulation of Bacteria-Gut-Brain Signals. Trends in Neurosciences, 39(11), 763–781. 10.1016/j.tins.2016.09.002

Seeman, M. V. (2021). The gut microbiome and antipsychotic treatment response. Behavioural Brain Research, 396, 112886. 10.1016/j.bbr.2020.112886

Spitzer, S. O., Tkacz, A., Savignac, H. M., Cooper, M., Giallourou, N., Mann, E. O., Bannerman, D. M., Swann, J. R., Anthony, D. C., Poole, P. S., & Burnet, P. W. J. (2021). Postnatal prebiotic supplementation in rats affects adult anxious behaviour, hippocampus, electrophysiology, metabolomics, and gut microbiota. iScience, 24(10), 103113. 10.1016/j.isci.2021.103113

Subramaniam, S., Elz, A., Wignall, A., Kamath, S., Ariaee, A., Hunter, A., Newblack, T., Wardill, H. R., Prestidge, C. A., & Joyce, P. (2023). Self-emulsifying drug delivery systems (SEDDS) disrupt the gut microbiota and trigger an intestinal inflammatory response in rats. International Journal of Pharmaceutics, 648, 123614. 10.1016/j.ijpharm.2023.123614

Subramaniam, S., Kamath, S., Ariaee, A., Prestidge, C., & Joyce, P. (2023). The impact of common pharmaceutical excipients on the gut microbiota. Expert Opinion on Drug Delivery, 1–18. 10.1080/17425247.2023.2223937

Tomizawa, Y., Kurokawa, S., Ishii, D., Miyaho, K., Ishii, C., Sanada, K., Fukuda, S., Mimura, M., & Kishimoto, T. (2021). Effects of Psychotropics on the Microbiome in Patients With Depression and Anxiety: Considerations in a Naturalistic Clinical Setting. International Journal of Neuropsychopharmacology, 24(2), 97–107. 10.1093/ijnp/pyaa070

Tuson, H. H., Foley, M. H., Koropatkin, N. M., & Biteen, J. S. (2018). The Starch Utilization System Assembles around Stationary Starch-Binding Proteins. Biophysical Journal, 115(2), 242–250. 10.1016/j.bpj.2017.12.015

Vacca, M., Celano, G., Calabrese, F. M., Portincasa, P., Gobbetti, M., & De Angelis, M. (2020). The Controversial Role of Human Gut Lachnospiraceae. Microorganisms, 8(4), 573. 10.3390/microorganisms8040573

Zafar, H., & Saier, M. H. (2021). Gut *Bacteroides* species in health and disease. Gut Microbes, 13(1), 1848158. 10.1080/19490976.2020.1848158

